# Sequence-function mapping of proline-rich antimicrobial peptides

**DOI:** 10.1101/2024.01.28.577586

**Authors:** Jonathan Collins, Adam McConnell, Zachary D. Schmitz, Benjamin J. Hackel

## Abstract

Antimicrobial peptides (AMPs) are essential elements of natural cellular combat and candidates as antibiotic therapy. Elevated function may be needed for robust physiological performance. Yet, both pure protein design and combinatorial library discovery are hindered by the complexity of antimicrobial activity. We applied a recently developed high-throughput technique, sequence-activity mapping of AMPs via depletion (SAMP-Dep), to proline-rich AMPs. Robust self-inhibition was achieved for metalnikowin 1 (Met) and apidaecin 1b (Api). SAMP-Dep exhibited high reproducibility with correlation coefficients 0.90 and 0.92, for Met and Api, respectively, between replicates and 0.99 and 0.96 for synonymous genetic variants. Sequence-activity maps were obtained via characterization of 26,000 and 34,000 mutants of Met and Api, respectively. Both AMPs exhibit similar mutational profiles including beneficial mutations at one terminus, the C-terminus for Met and N-terminus for Api, which is consistent with their opposite binding orientations in the ribosome. While Met and Api reside with the family of proline-rich AMPs, different proline sites exhibit substantially different mutational tolerance. Within the PRP motif, proline mutation eliminates activity, whereas non-PRP prolines readily tolerate mutation. Homologous mutations are more tolerated, particularly at alternating sites on one ‘face’ of the peptide. Important and consistent epistasis was observed following the PRP domain within the segment that extends into the ribosomal exit tunnel for both peptides. Variants identified from the SAMP-Dep platform were produced and exposed toward Gram-negative species exogenously, showing either increased potency or specificity for strains tested. In addition to mapping sequence-activity space for fundamental insight and therapeutic engineering, the results advance the robustness of the SAMP-Dep platform for activity characterization.

## Introduction

Antimicrobial peptides (AMPs) are a compelling source of antibiotic therapeutics.^1^ Yet improved potency, selectivity, and stability could aid physiological performance.^2^ Understanding sequence-function relationships would identify better molecules, empower further engineering, and advance fundamental understanding of this important class of molecules. Sequence-function mapping has been highly effective for fundamental study and functional engineering across many proteins.^3,4^ Yet the complexity of antimicrobial activity typically hinders the high-throughput evaluation needed to elucidate the immense, rugged sequence-function landscape.

Recently, several technology platforms have been developed for library-scale evaluation of AMP activity. In two instances, a cellular host produces an AMP variant in a format that enables the AMP to kill and/or inhibit growth of the cellular host. In a library of cells, each with distinct variants, potent variants will be reduced in relative abundance whereas non-potent variants will propagate. Deep sequencing pre- and post-expression of AMPs enables identification of relative potencies. In one such approach, surface-localized antimicrobial display (SLAY)^5^, AMPs are tethered to the extracellular surface via a polypeptide linker to the transmembrane protein OmpA. AMP-OmpA fusions can then act on extracellularly-accessible targets. In another approach, sequence-activity mapping of AMPs via depletion (SAMP-Dep)^6^, AMPs are expressed intracellularly, thereby enabling assessment of intracellular potency through population dynamics at varying induction/expression levels (Figure 1). This method was developed and validated using oncocin, resulting in elucidation of the sequence-activity landscape and identification of variants with elevated potency.^6^ It was later applied, with the addition of periplasmic transport, to study endolysins that act upon peptidoglycans.^7^ Herein, the utility of SAMP-Dep was extended to additional proline-rich AMPs to garner greater information in developing potential therapeutic antibiotic candidates from this protein family.

**Figure 1.**
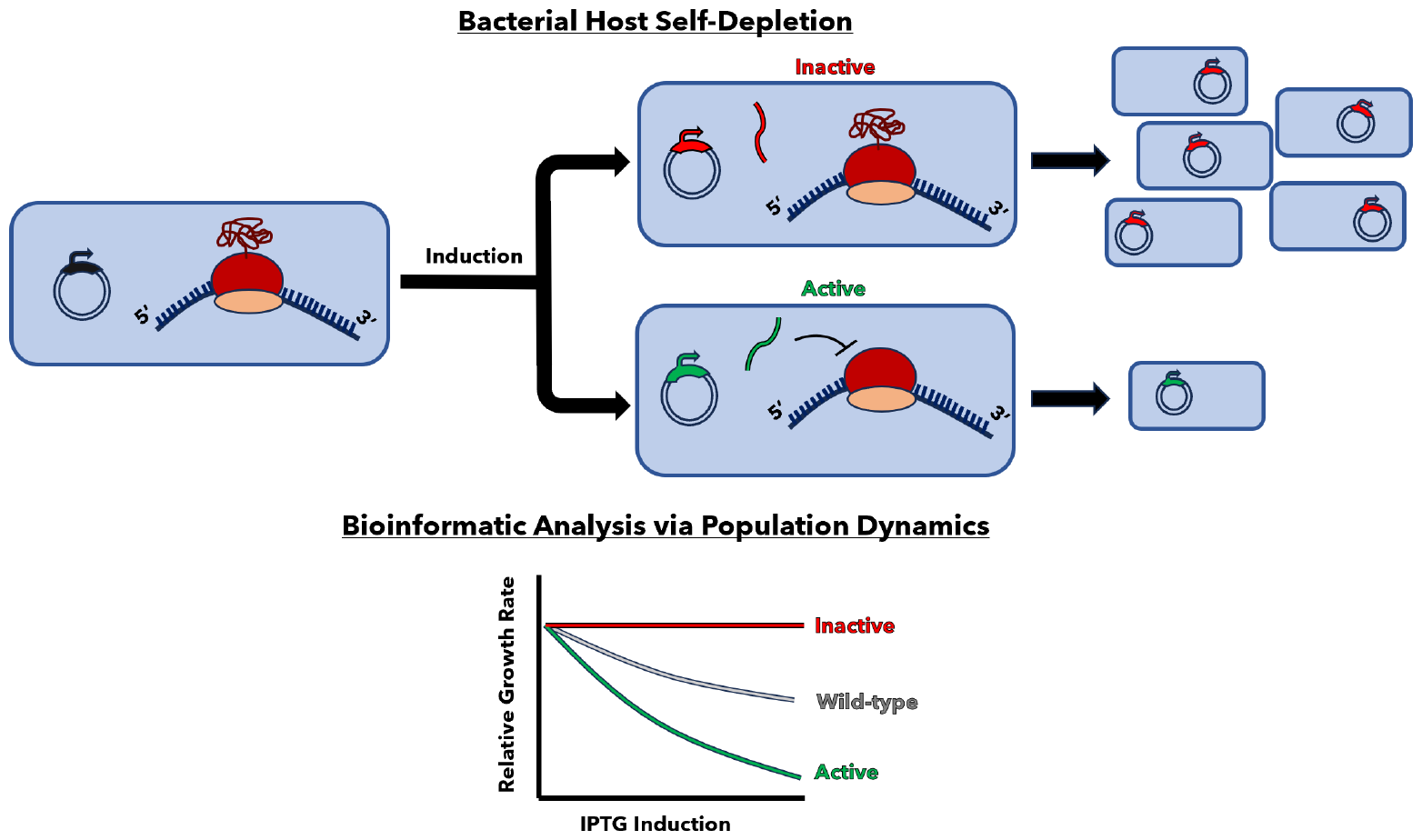
Outline of the SAMP-Dep platform steps. Bacterial hosts (*E. coli* LysY/I^q^) are transformed with an expression vector encoding PrAMP variants. Hosts are induced to generate internal expression of encoded PrAMPs and allowed to propagate. Inactive PrAMPs do not cause ribosomal inhibition, leaving growth unaffected. Active PrAMPs decrease reproduction of their host proportionally to their level of potency and expression. DNA is harvested from samples before and after incubation to determine relative growth rate changes of each DNA variant over the entire population. The relative changes in growth rate are then analyzed by DNA variant counts via deep sequencing. Greater negativity in the slope of growth rate versus inducer concentration equates to greater potency.

Proline-rich AMPs are widely found in nature from insects (*e*.*g*. metalnikowin 1 (Met) in fruit flies and apidaecin 1b (Api) in honeybees^8^) to mammals (Tur1a in bottlenose dolphins^9^).^10^ Proline-rich AMPs are non-lytic peptides that act mainly through stereo-specific inhibition of protein translation via the 70S ribosome within Gram-negative bacteria.^8,11^ These peptides were initially thought to mainly inhibit DnaK, an important protein chaperone; however experiments with *E. coli* unable to express DnaK only showed slight differences in activity. Moreover, co-immunoprecipitation identified high ribosomal binding, highlighting the main target as the 70S ribosome for proline-rich AMP activity.^12–14^ Several characteristics define the family of proline-rich AMPs. First, they tend to have an overall positive charge that is hypothesized to aid in membrane interaction and internalization.^15^ Consistent with their name, they have a high percentage of proline (>25%).^10^ This creates a hybrid random coil/poly-proline helical structure hypothesized to allow insertion into the ribosome (Figure 2), since this hybrid helix has a smaller width and extended structure compared to a standard α-helix.^16^ Almost ubiquitously, members incorporate at least one PRP motif that is flanked by a tyrosine residue (commonly a YR/YL segment) hypothesized to be the main functional motif within these peptides.^17^ Further, in all resolved peptide-ribosome complexes the PRP motif is in the exact same orientation and position, supporting this hypothesis.^15^

**Figure 2.**
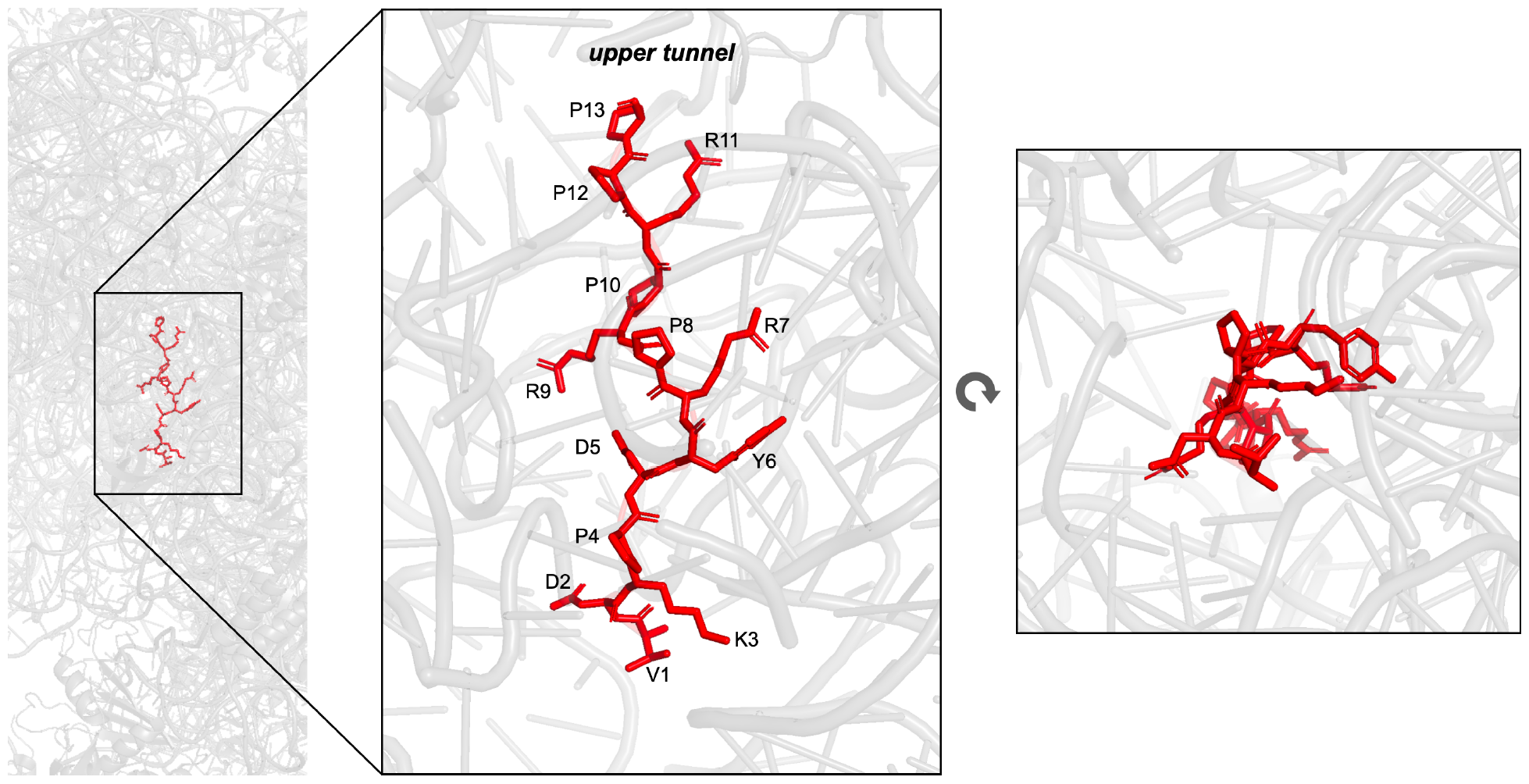
PrAMP/ribosome structure. The first 13 residues of Met are resolved in complex with the ribosome (PDB: 5HCP).

In the current study, we used SAMP-Dep to map the sequence-activity landscapes for Api and Met. High reproducibility and alignment with existing hypotheses/studies further validated the SAMP-Dep platform, extending it across members of the PrAMP family of peptides. Several PrAMPs behave similarly in mutational tolerance and homologous residue performance including proline mutational tolerance, which is highly site-specific rather than density dependent, with required maintenance within the PRP domain. Epistatic interactions between residues following the PRP domain and extending into the ribosomal exit tunnels appeared to be the most important and consistent. Exogenous exposure of externally synthesized PrAMP variants validated activity identified from internal production and exposure. The results provide sequence-activity maps for two important AMPs and further the utility of the SAMP-Dep engineering platform.

## Results

### SAMP-Dep platform and AMP diversification

Six proline-rich AMPs were initially selected based on their distinctive chemical diversity and ribosome binding mechanics within the proline-rich AMP family^10^ (Supp Fig 1). The abilities of these AMPs to deplete their host bacterium upon induction of protein expression, which is requisite for the SAMP-Dep concept, was assessed. The genes for each AMP were cloned into a pET expression plasmid with a T7lac promoter system and transformed into T7 LysY/Iq *E. coli*. Induction with isopropyl ß-D-1-thiogalactopyranoside (IPTG) dramatically slowed growth for Met, Api, and Tur1a (Figure 3). Three other proline-rich AMPs tested (arasin 1, heliocin, and pyrrhocoricin) were inactive. Conversely to the three actives, all of which have a single independent PRP motif, the inactive AMPs have none (heliocin) or two (arasin and pyrrhocoricin). Since the PRP motif is hypothesized to be necessary for DnaK and ribosome binding, heliocin lacking this motif may indicate it behaves differently than other proline-rich AMPs. Neither of arasin 1’s two PRP motifs are in proximity to a tyrosine and pyrrhocoricin requires T17 glycosylation for high functionality^8^, which likely contributes to their lack of self-depletion within the platform. Further, arasin 1’s intracellular target has not been elucidated meaning its mechanism of action is still not fully understood and may not be present in the engineered/model organism used in this study^18,19^.

**Figure 3.**
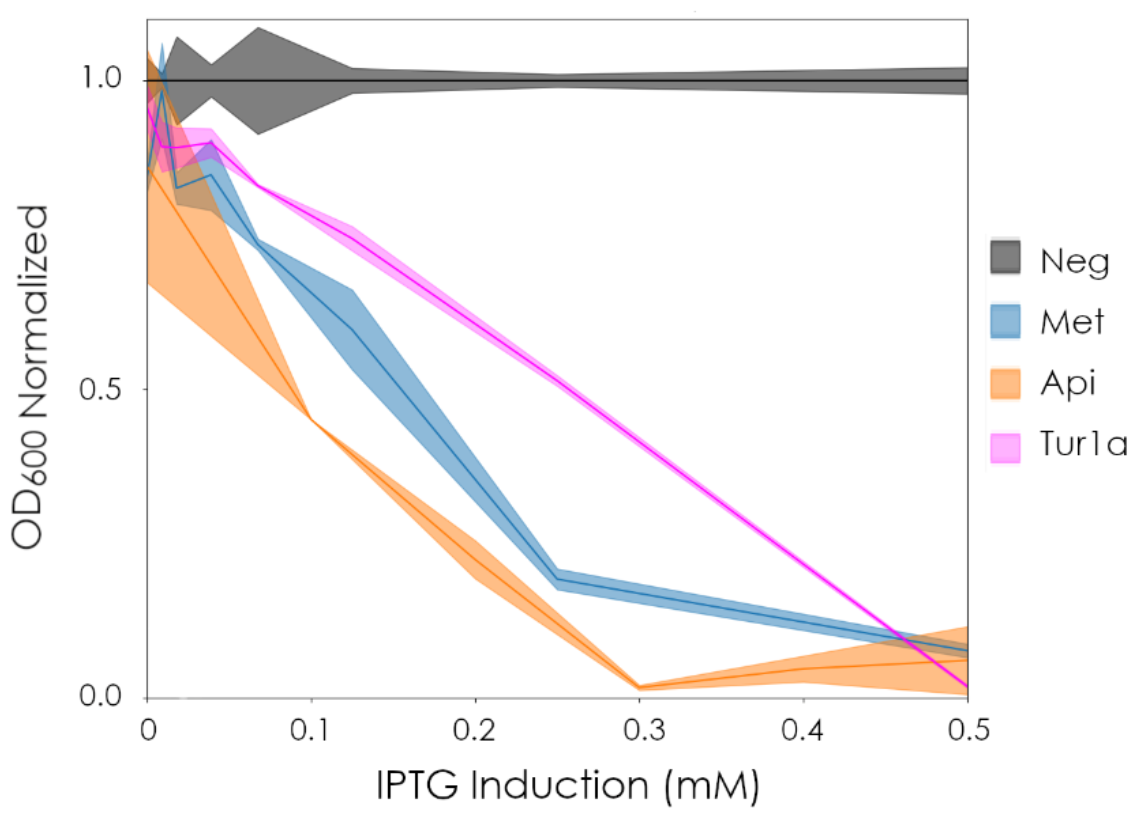
Induction of AMP expression hinders host growth. Bacterial growth upon induction of expression of wild-type Met, Api, Tur1a, or negative control with prematurely stopped AMP gene normalized to the negative control without AMP gene.

To compare and contrast our assembled dataset relative to the oncocin dataset^6^, we opted to evaluate the two active AMPs with the most homology with oncocin. Met exhibits identity to oncocin at 11 sites with four mutations and truncation of the final four amino acids (Table 1). Api shares the PRP motif of oncocin – and more broadly a YIPQPRPP motif homologous to oncocin’s YLPRPRPP motif – but has reduced homology elsewhere. Conversely, Tur1a possesses the PRPxRR motif and a YLPRP motif but otherwise bears little resemblance to oncocin; thus it will be the subject of a separate study. Met is a class I proline-rich AMP like oncocin; Api is the only known class II proline-rich AMP binding in the reverse orientation within the ribosome. This change in binding mechanics changes the overall function of the peptide from an initial elongation inhibitor to a termination inhibitor.^9,15,20–23^ This may additionally presume Api’s PRPPH motif is acting as the main binding/inhibition segment, only in reverse.

**Table 1.**
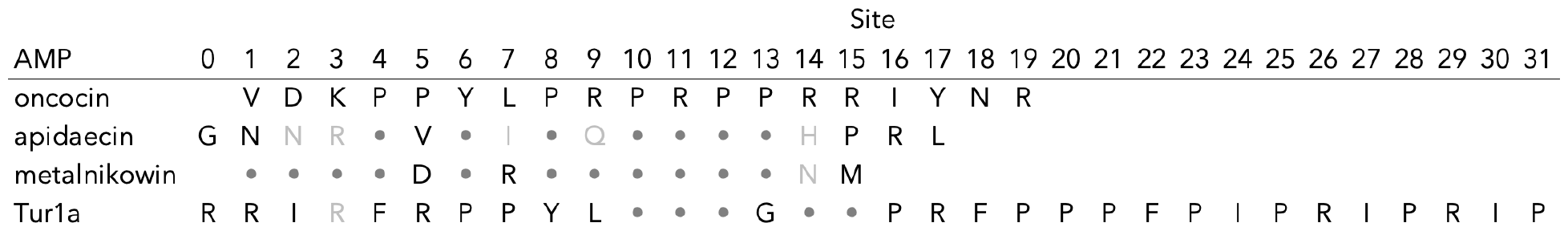
AMP sequences aligned to oncocin. Amino acids identical to oncocin are indicated as •. Amino acids homologous (BLOSUM62 ≥0) to oncocin residues are indicated in gray.

Doublet saturation mutagenesis libraries were assembled for both AMPs in i / i+1 and i / i+2 configurations in a sliding window through each AMP. Gene libraries were assembled from degenerate oligonucleotides encoding for all 4×10^2^ protein variants (and 1×10^3^ DNA variants) at each set of doublets for a total of 27,648 Met variants and 33,792 Api variants in hypothetical diversity. Bacterial transformation yielded 9 ± 3 million transformants (675 ± 185-fold oversampling of sequence space) for DNA preparation and 33 ± 9 million transformants (1200 ± 340-fold) for host creation for producing Met variants. For Api, bacterial transformation yielded 15 ± 1 million transformants (860-fold) for DNA preparation and 30 ± 3 million transformants (880 ± 95-fold) for host creation. DNA sequencing indicates that 93% of the designed Met library and >99% of the Api library were present in all replicates at uninduced and initial conditions. Further, numerous unintended mutants arose during library synthesis (Supp Fig 2).

### Sequence-function mapping via SAMP-Dep

Libraries for the two AMPs, across four independent replicate transformations for each, were grown and induced to express each AMP. The population was deep sequenced before and after several levels of induction strength. The slope of clonal frequency in the population versus inducer strength was computed for each genetic variant and resultant protein sequence. Clonal slopes were reproducible across quadruplicate runs (average inter-run correlation coefficient, ρ = 0.76; Figure 4A), with increasing correlation strength for clones with higher initial read depth (Figure 4A) (as low initial frequencies have higher uncertainty as well as less available range for depletion). Importantly, 79% of all clones have at least 20 reads in all uninduced replicates, which corresponds to ρ = 0.90 on average for Met (Figure 4A,B). For Api, these metrics are 84% and ρ = 0.92 (Figure 4A,B). This 20 read threshold was used to filter sequences represented enough in each replicate to inform analysis within the rest of the study. Assay validity is further supported by strong correlation (ρ = 0.99 for Met and ρ = 0.96 for Api (Figure 4C)) across synonymous genetic variants as well as the lack of activity of mutants with early truncations (see Figure 5B,C). 54% of variants in Met and 57% in Api exhibit no activity (all replicate slopes not significantly different than zero; Figure 5A), which highlights the sensitivity to mutation within these small peptides. Yet, numerous tolerated and beneficial mutations were also observed; 3% of variants for Met and 17% of variants for Api exhibit activity statistically superior to wildtype showing the potential for improved potency.

**Figure 4.**
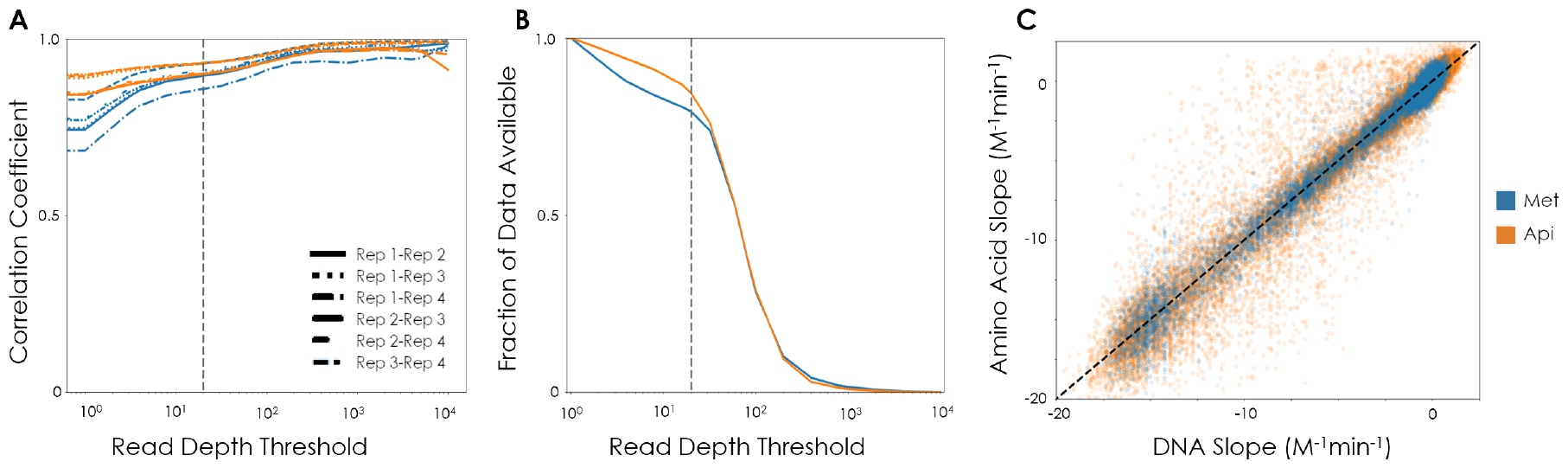
SAMP-Dep data exhibit high reproducibility. (A) The replicate-to-replicate correlation is plotted for all variants at or above the indicated number of clonal reads in the 0 mM sample. (B) Distribution function indicating the fraction of protein variants with at least the indicated number of clonal reads in the 0 mM sample, dashed line indicates the 20 read threshold used for analysis on data. (C) The slope of frequency versus inducer concentration for each protein variant in each replicate is plotted against the slope of the corresponding gene, dashed line represents when the Amino Acid Slope = DNA Slope.

**Figure 5.**
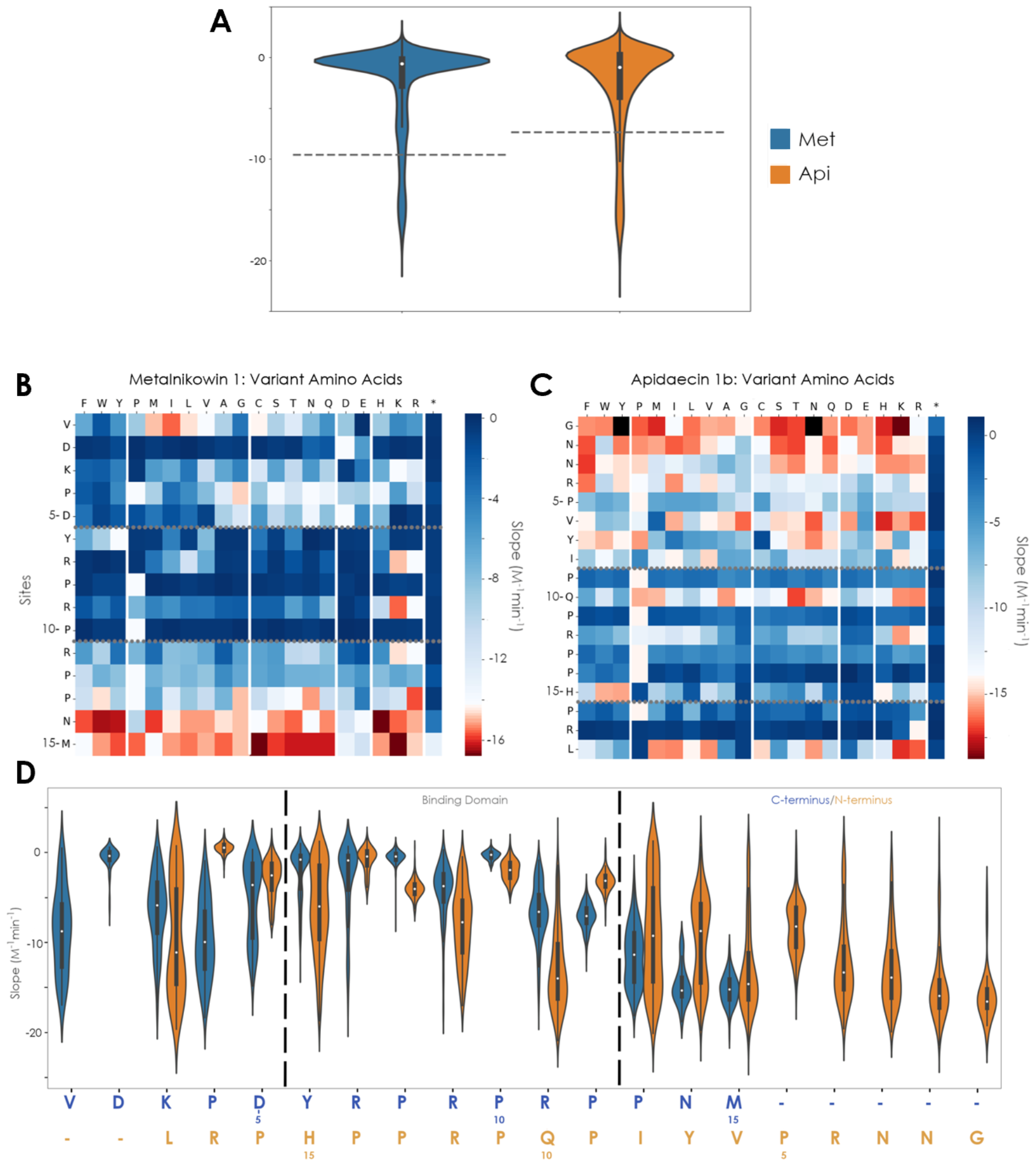
AMP sequence-function maps of each peptide. (A) Slope distributions for each AMP. Heatmaps: each square represents the average slope (clonal frequency versus inducer concentration) of all variants with that column’s amino acid at that row’s site for (B) Met and (C) Api. Black is no data. Wild-type baseline in graphs is average wild-type variant slope. (D) Mutational tolerance of both PrAMPs aligned around the binding/inhibition domain residues.

Overall, beneficial mutations in Met dominantly occur near the C-terminus (Figure 5B). A few exceptions are the homologous mutations of V1(I, M, or L) and R-to-K mutations at sites 7, 9, and 11. The mutational tolerance for Api has similarities and contrasts. Similarly, many of the most beneficial mutations reside within the three terminal sites, in Api’s case the N-terminus (Figure 5C). These observations are hypothesized due to its reverse binding orientation within the ribosome, where its N–terminal mutational tolerance reflects that of the C–terminus of the screened class I PrAMPs. Conversely to Met, Api exhibits a broader array of sites with beneficial mutations, extending out to the first seven sites at the N-terminal end as well as Q10 and L18 (both to multiple alternate amino acids), R12K, and H15 (to either alternate hydrophilic aromatic). This aromatic-only site (15) reverse aligns with similar behavior at Y6 in Met (Figure 5D). Similarly, oncocin exhibits Y6H mutational tolerance and numerous beneficial C-terminal mutations.^6^

Analysis of site-wise mutational impacts reveals multiple intolerant sites in both peptides (Figure 5B,C). Four sites relatively intolerant to mutation in Met are D2, Y6, P8, and P10. D2 and Y6 maintain some activity with chemically similar residues; however, the first two prolines (sites 8 and 10) in the overlapping PRP motif exhibit no activity upon mutation of either. Unlike the two prolines of the PRP motif, the central R is moderately tolerant of mutation. This R benefits from mutation to the other strong cation, K. Moderate activity is enabled by mutation to H, Q, or P and it somewhat tolerates polar uncharged residues T and Q, but loses significant activity upon other mutations including loss of all activity upon charge switch to D or E. The following arginine and proline within the overlapping motif are more amenable to mutations but still generally result in decreased potency of the molecule. The KPD motif at sites 3-5 maintains activity upon mutation to an array of alternatives. In addition, the final amino acid can be deleted with no appreciable detriment to activity, aligning with the exact site oncocin begins to allow truncations without decreased potency.^6^ For Api, site R17 is intolerant to mutation similarly to the second site from the terminus (D2) in Met. P9, P11, P13, and P14 within the binding/inhibition domain also significantly lose potency upon substitution to other amino acids (see below for more detail on proline mutations). The central arginine of the PRP motif also only benefits from strongly cationic lysine and tolerates hydrophobes P, M, I, and L, but substantially loses potency upon charge switching to anionic residues D and E. Adjacent to the PRP motif, Q10/P9/I8 exhibits a similar tolerance profile as the corresponding RPP motif in Met. Cationic mutations at Q10 increase potency, allowing creation of an overlapping PRP domain in Api. Lastly, no early truncations maintain any activity; however, there is a sequence including a full N-terminal codon shift removing the first residue present with three other point mutations causing potency significantly better than wild-type (YPKPVYIPQPRPPHPRL, slope: -15 M^-1^min^-1^).

### Mutational Relationship to Homology

We next evaluated sequence-activity relationships from a global perspective. Mutant activity correlates with the homology (as determined from BLOSUM62 score^24^) of the wild-type and mutant amino acids (Figure 6A), which aligns with prior work^25^. Api exhibits more mutational tolerance than Met for all levels of mutational homology, except for highly non-homologous mutations in which both perform poorly. Notably, highly homologous mutations have a slightly worse performance than moderately homologous mutations for Met mainly driven by other aromatics substitutions to Y6 that do not perform well.

**Figure 6.**
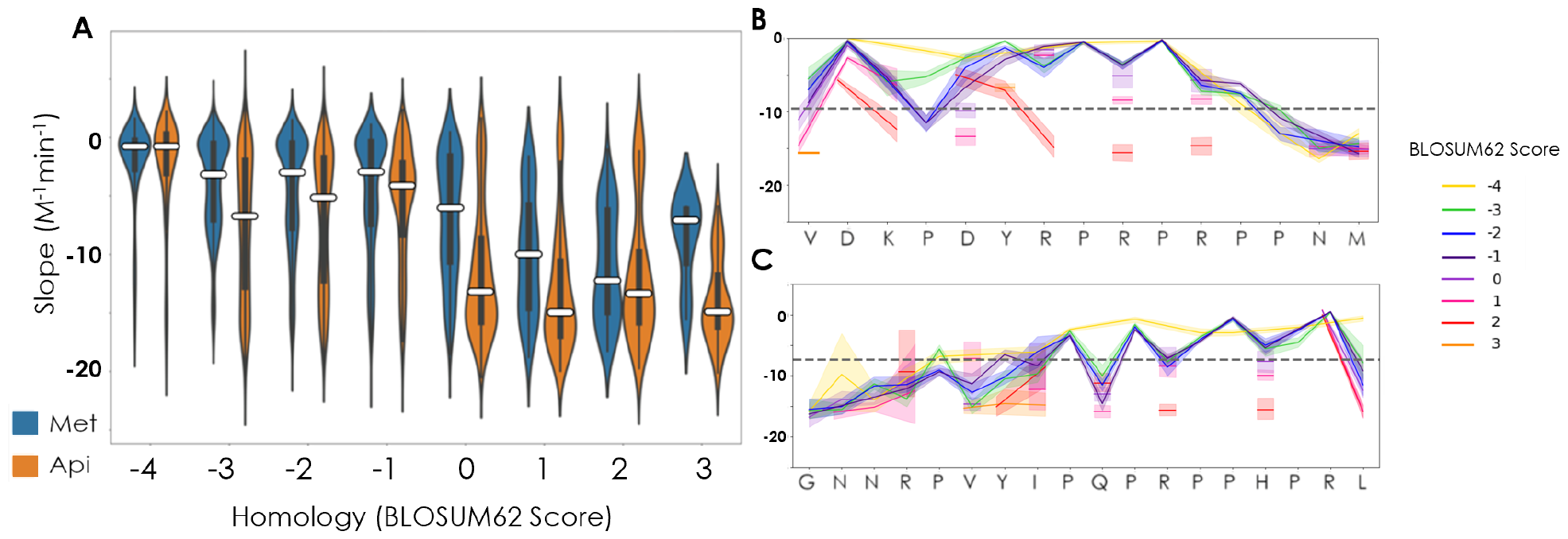
Homology benefits to potency are highly site/region specific. A) distribution of all slopes with single mutations scored with BLOSUM62 matrix for both Met and Api. (B, C) Lineplots showing mutations at each BLOSUM62 score for Met (B) and Api (C).

The impact of homology on mutational tolerance is spatially dependent. Along alternating sites in Met (1, 3, 5, 7, 9, 11, and 15) on one ‘face’ of the peptide, activity can be achieved with highly homologous mutations (Figure 6B). The C-terminus is the only section of the peptide allowing highly non-homologous mutations. Mirrored in Api, highly non-homologous mutations are primarily accepted at the N-terminus, as well as Q10 and L18. Similarly, there is a striking alternating pattern of tolerance directly related to the proline residues offering the least amount of mutational tolerance besides P4 in Met and P5 in Api. These domains align very well between the two peptides in their mutational tolerance and residue type, alluding to similar functional mechanics in ribosome inhibition. This dependency on homology is highest within the binding/inhibition domain. In both peptides, residues in the binding domain only allow highly homologous mutations.

### Functional Effects of Proline

Prolines comprise 33-38% of these peptides and are hypothesized to be a necessary structural feature in binding/inhibition through destabilizing an α-helix from forming and extending contacts to rRNA in the main ribosomal subunit.^10,16^ Proline mutational tolerance varies by AMP and site. For Met, 100% of proline mutants within the PRP motif have slopes less than wild-type as compared to 58% of other proline mutants and 79% for non-proline mutants (Figure 5B-D). The PRP motif mutational impacts are not only broad, but they are substantial, as 100% reduce slope two-fold from wild-type as compared to only 56% for non-proline mutants. In Api, proline mutational impact is broader with 100% of proline mutants within the PRP motif, and 74% of non-motif prolines, having slopes less than wild-type as compared to 53% for non-proline mutants. Again these are more likely to be significantly detrimental, as 76% in the PRP motif reduce slope at least two-fold from wild-type compared to only 43% for non-PRP prolines and non-proline mutants.

Improved activity with reduced proline density is achieved primarily via mutation away from P13 or P4 in Met and P5 in Api (Figure 7). Mutation of P13 and P16 in Api are generally deleterious, but activity can be recovered or increased with select simultaneous local mutations (Figures 7, 8). For Met, only one sequence performed significantly better while removing two wild-type prolines: the undesigned triple mutant P4T, P13R, M15W. For Api, no sequences with two localized wild-type prolines removed were able to perform better than the wild-type baseline sequence. Elevated activity with increased proline density is achieved via highly different scenarios between the peptides. Site 15 is the only location to add a proline to boost activity for Met, whereas in Api a single or double proline addition can boost activity at all sites in or before the binding domain.

**Figure 7.**
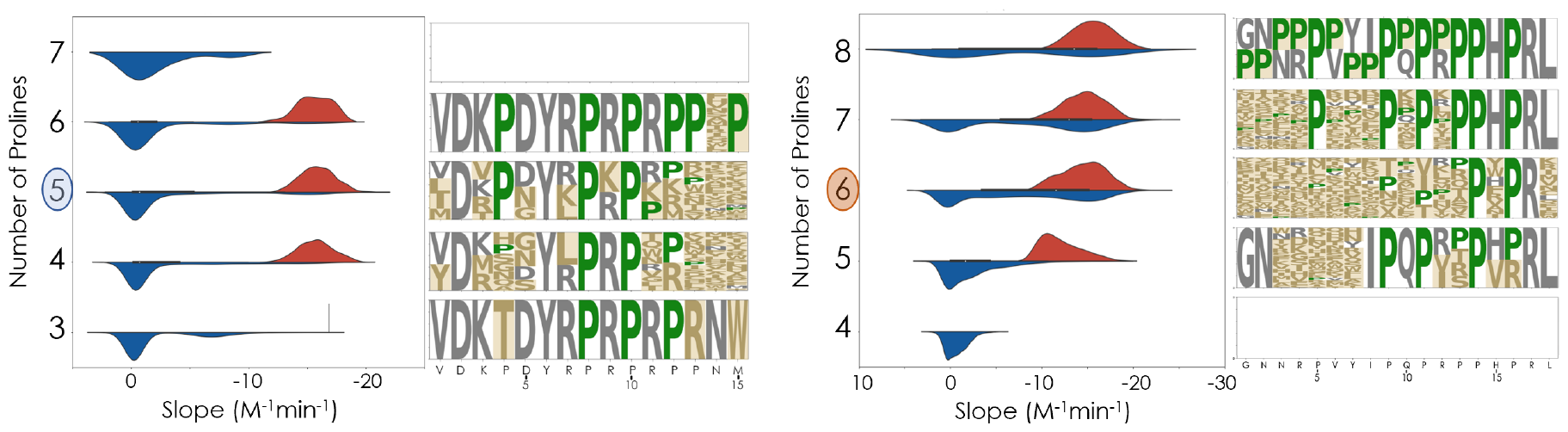
The impact of proline on AMP activity. The distribution of slopes (clonal frequency versus inducer concentration) across all variants with the indicated number of prolines is presented. The wild-type proline count is highlighted on the axis. Blue violin plots correspond to all variants. Red plots correspond to variants significantly better than wild-type. On the right of the plots are sequence logos of the significantly better sequences (residue sizes are weighted by their average slope difference from wild-type baseline). Orange highlighted sections indicate mutations. Prolines colored green.

**Figure 8.**
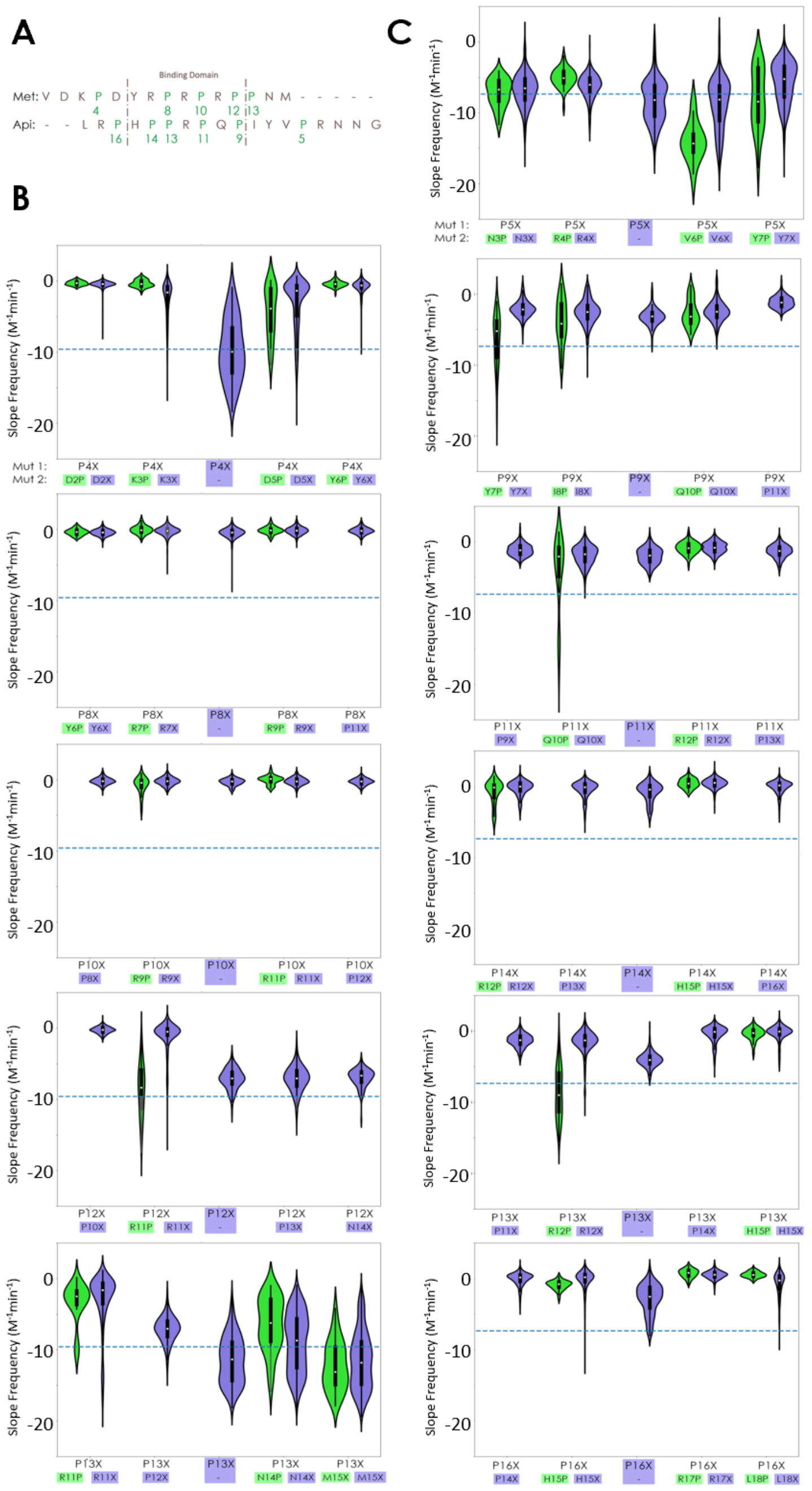
The impact of proline shifting on AMP activity. The distribution of slopes corresponding to local mutations around substituted proline. Green violins represent shifted prolines. Blue-violet distributions represent all other multimutants. A) shows aligned sequences with wild-type prolines highlighted, B) represents data for Met, and C) represents data for Api.

The library design also permits proline shifting, i.e. wild-type proline substitutions along with a simultaneous proline addition within ± 2 residues. Within this frame, Api is slightly more amenable to proline shifting than Met with 18% of proline shifts as active or more active than the wild-type peptide compared to 12% for Met (Figure 8). With the highly tolerant prolines P4 and P13 in Met, both can shift to sites 5 or 14/15, respectively. However, these do not necessarily perform better than other multimutants. Beyond these highly tolerant sites, P12 can shift to site 11 (R11P, P12X) to recover P12 mutational deficits and in some cases increase potency significantly better than other multimutants. P8 and P10, again, are not amenable to mutation and cannot recover wild-type activity by shifting to another local site. In Api, several key sites have a high likelihood of increasing activity with shifted prolines. As mentioned previously, P5 is generally tolerant of mutations but shifting to site 6 (P5X, V6P) almost always increases potency and performs significantly better than other multimutants (Figure 8). To a lesser degree, increases in activity can also be seen in shifting to site 7 (P5X, Y7P). This Y7 site also has an ability to shift with the site 9 proline to recover and increase activity (Y7P, P9X) greater than other multimutants. Site 8 shifting (I8P, P9X) to a lesser degree can recover activity, but not significantly better than other multimutants. P11 and P13 are very intolerant to single mutations, but shifting to sites 10 and 12, respectively, (Q10P, P11X and R12P, P13X) can recover and increase activity more likely than other multimutants. Lastly, P14 and P16 mutations mainly decrease potency and are unable to recover wild-type activity when locally shifted.

### Epistatic Analysis for Multimutants

To generalize beyond proline shifting to all dual mutation options, we evaluated epistasis to identify double-mutants with a change in slope more substantial than expected from additivity of the two single mutations. For Met, sites 11 and 12 as well as R11 and P13 had notable interactive effects (Figure 9). When R11 was mutated to either proline or a branched hydrophobic residue (I/V) and P12 was mutated to a cationic residue (K/R/H), it resulted in a potent double mutant superior to the additivity of the single mutations. When R11 is mutated to either D or H and P13 is mutated to a cationic residue (K/R), positive epistasis was also observed. Strikingly, many other double mutations at these residues (11+12 or 11+13) result in negative epistasis yielding an inactive variant, further demonstrating the interaction between these sites. Lastly, a beneficial interaction is also achieved when proline is shifted from site 4 to site 5 with site 4 flipping local charge to a cationic residue (K/R) versus the original D5.

**Figure 9.**
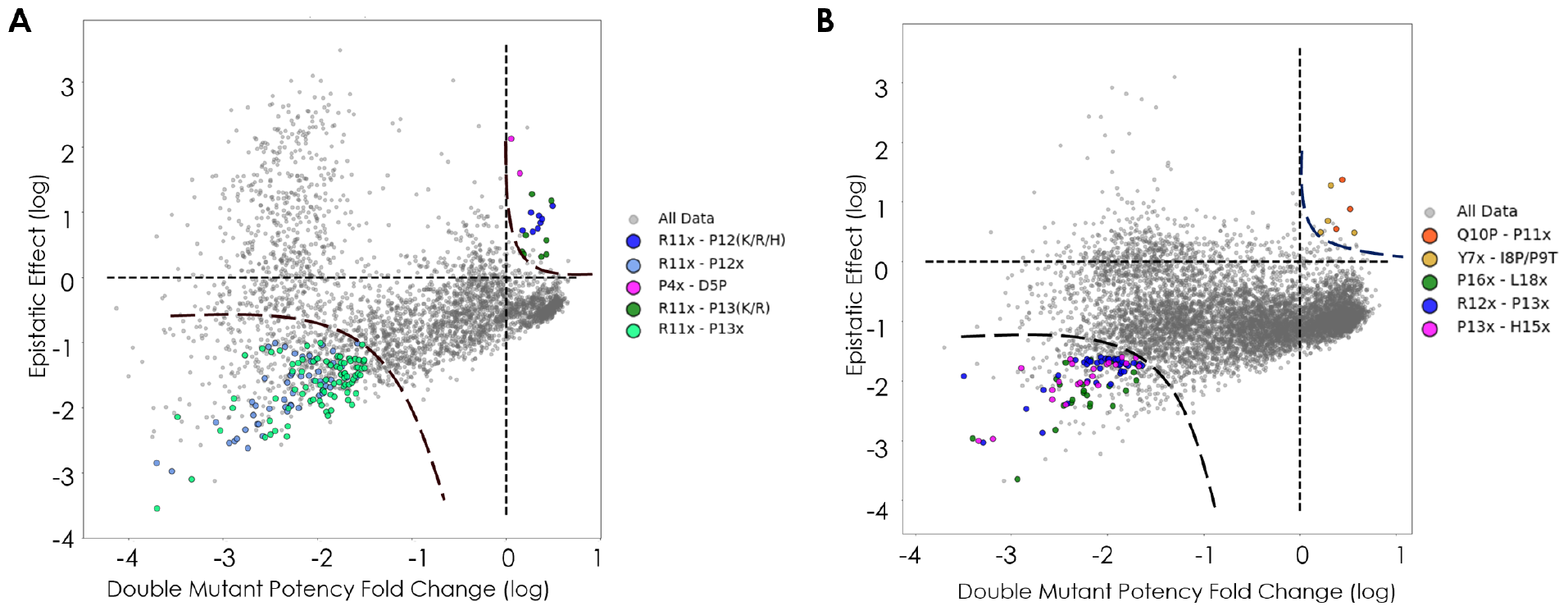
Residue interaction within peptides. Epistatic terms between two subsequent mutations given their SAMP-Dep double mutation performance, calculated with average slope of all amino acid sequence variants for A) Met and B) Api. Important and consistent residue interaction points highlighted for each peptide.

Within Api, the most frequent form of positive epistasis yielding more potency than wild-type was observed in a proline shifted from site 11 to Q10P and site 11 hosting a branched hydrophobic residue (V/I/T). Y7 has several positively epistatic interactions: in concert with I8P, a mutation to a branched hydrophobic residue at Y7 (V/I) increases the dual mutant performance; if the proline mutation instead occurs as Y7P, then P9T increases the dual mutant performance. The negatively epistatic interactions yielding inactivity include binding domain residue combinations of (R12, P13) and (P13, H15). These are highly important residues, where single mutations that are somewhat tolerated cannot compensate with another simultaneous mutation within this segment. Lastly, L18 can also tolerate several mutations very well (some even increasing performance) but does not have the same effect when subsequent mutations happen at P16.

### Extracellular Treatment with Selected Variants

We selected several variants (Figure 10A) for extracellular treatment based on their slope distribution over all unique DNA sequences. The top three peptides with greatest negative averaged DNA slopes were taken and produced through solid phase synthesis and exposed to cultured bacteria to assess growth inhibition. The most statistically significant inactive sequence and the wild-type sequence were used as controls (Figure 10B). Another sequence was chosen as a “special case” based on interesting proline contents paired with high SAMP-Dep potency.

**Figure 10.**
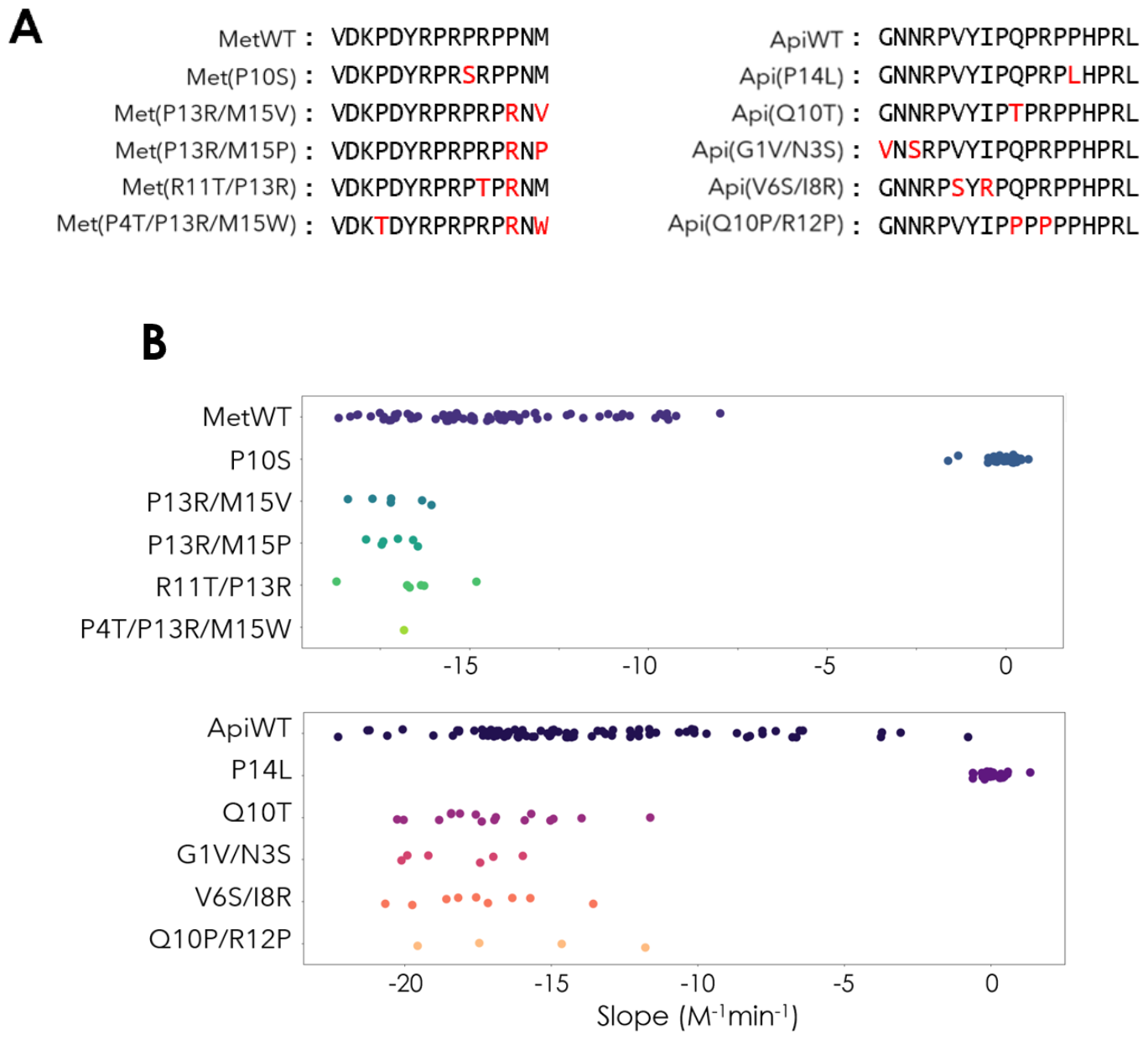
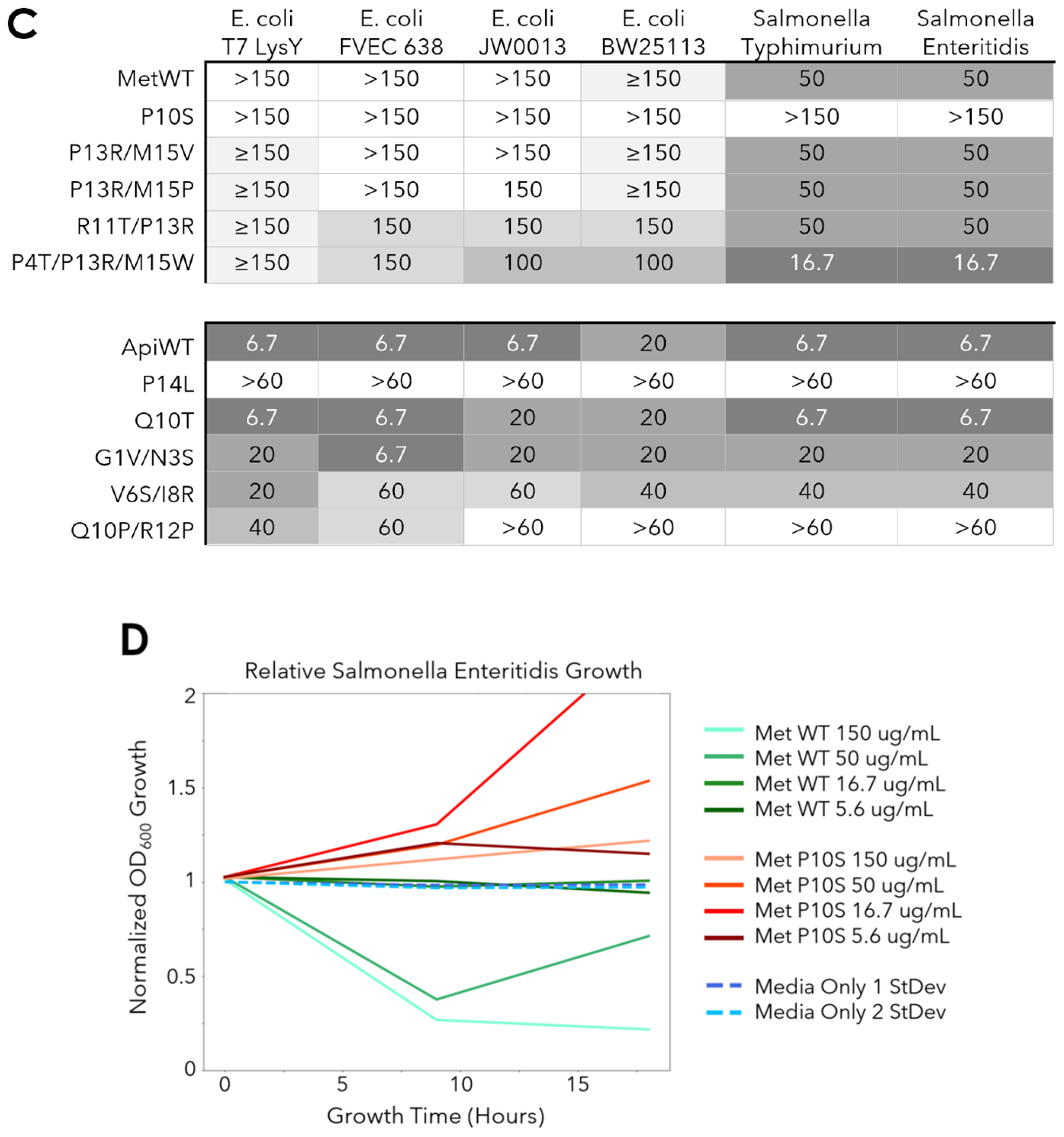
Variants identified for external exposure in susceptible species from SAMP-Dep Assay. Peptide sequences selected are outlined in A). These include wild-type variants, the most statistically inactive variant, the three most statistically active variants, and a special active case with SAMP-Dep performance in B) for Met and Api. C) shows the performance against six strains of two species mirroring activity assays shown in Lai (2019)^26^. Concentrations shown are listed in µg/mL for all peptides and strains.

Wild-type Met exhibited very little potency for *E. coli* but did have activity for the tested *S. enterica* species (Figure 10C). In contrast, Api had high potency for almost all species tested, slightly decreasing for *E. coli* BW25113. Both internal negative controls displayed no inhibitory activity in any species. The most potent variant for Met was the undesigned multimutant, P4T/P13R/M15W; this was the only sequence with 2 prolines removed to maintain internal activity. It increased activity toward *S. enterica* species three-fold and had the highest potency for *E. coli* FVEC 638, JW0013, and BW25113. The remaining variants maintained comparable activity to the wild-type Met peptide. Regarding Api variants, the three most active mutants in the intracellular SAMP-Dep context exhibited potency upon extracellular treatment. Q10T is the most comparable in potency to wild-type, only decreasing potency for *E. coli* JW0013, which is a DnaK knockout strain. Interestingly, the G1V/N3S variant exhibits mild specificity for *E. coli* FVEC 638 strain, decreasing in activity toward all others tested. The least potent yet still active variant is Q10P/R12P, which maintains internal activity but creates a segment of six straight prolines within the hypothesized binding domain. This variant maintains its highest potency for the T7 LysY *E. coli* in an exogenous context reflective of the platform in which it was identified, but significantly reduces potency for all other strains.

## Discussion

Thorough mutational scanning of AMPs^5–7,27^ has started to build our understanding of their sequence-function relationships, which elucidates fundamental mechanisms and empowers engineering toward therapeutic utility of these antibiotic candidates. We applied the SAMP-Dep platform to quantify the sequence-activity landscape of Met and Api. Mutations with the greatest benefits appeared within the C-terminal region of Met (Class I) and N-terminal region of Api (Class II). Previously, variances within the N-terminal region of apidaecins have correlated with changes to the range of susceptible species of the molecule.^8^ Therefore, beneficial mutations found within the N-terminus of Api and the C-terminus of class I proline-rich AMPs may coincide with increased specificity for the *E. coli* 70S ribosome being screened. To further consider structural connections to the elucidated landscape, it is worth noting that this highly variable region has been structurally resolved within the upper chamber of the ribosomal exit tunnel, which may have greater flexibility than the rest of the tunnel. This spatial region also only forms one hydrogen bond and single stacking pair between the ribosome and the wild-type C-terminal residues, which gives the potential for drastic binding improvements upon residue substitution^9,15,21,23^. Lastly, other ribosomal targeting constructs like macrolides have shown that specific amino acid sequences can still be produced while inhibitors are interacting with the exit tunnel^28,29^. Therefore, these beneficial mutations may either drive greater PrAMP-ribosomal interactions within this region of higher plasticity and fewer contacts or could also differentially restrict amino acid sequences capable of slipping past the inhibitor.

Additionally, in both peptides the hypothesized binding domain has very low tolerance to mutational changes with either single or multiple simultaneous mutations. The most important residues consist of the aromatic residue (site Y6 for Met, H15 for Api) and the motif’s proline residues (P8 and P10 in Met, P11 and P13/P14 in Api). However, the extended portion of an overlapping PRP domain in Met has much greater mutational tolerance than the rest of this segment. Certain prolines are able to be shifted within each peptide to either recover single mutational activity losses or even increase potency for the molecule. This suggests certain prolines are necessary for binding within the ribosomal A-site binding pocket or exit tunnel and others are structurally helpful in disrupting alpha-helix formation, elongating the peptide within the ribosome to maximize contacts^16^. These insights and the means by which they were procured provide a necessary and solid stepping-stone for developing PrAMPs into potent therapeutic agents.

We have effectively identified more potent and specific variants of Met and Api, with the potential to treat gram-negative bacterial infections. Previously, the only recorded inhibition of Met was seen at 62.5 µM in *E. coli* SQ110 (LPTD) strain that is highly sensitive to antibiotic activity^21^, no perceptive activity at 256 µg/mL for all tested species. The variants tested showed comparable to drastic increases for Met potency, especially for commonly pathogenic *S. enterica* strains. With Api variants, the wild-type peptide proved to be the most widely potent. However, select variants displayed certain specificity changes that could prove useful in selective treatments or pathways of evolutionary escape from susceptibility. Overall, we were only able to test a handful of identified active variants within an environmental exposure context but found variants able to generate or maintain high potency towards Gram-negative bacteria. Broader evaluation is needed to understand the differences between internal exposure and the peptide characteristics needed for internalization.

We have built on previous studies of oncocin^6^ and endolysins^7^ to further validate and refine the SAMP-Dep platform. Although deep sequencing drastically improves screening capacity, read depth limits throughput efficiency. Maximum calculable slopes show that an excess of 100 reads per sequence within each replicate-condition are needed for a sequence’s potency to robustly surpass the most active wild-type sequence (Supp Figure 3). Considering the distribution of sequence reads (Figure 4B), the number of genetic variants must be reasonably constrained with currently accessible sequencing resources, despite their notable efficiency. Nevertheless, the *n*/*n*+1 and *n*/*n*+2 libraries of 26,000 and 34,000 variants in the current work represent a substantial advance past earlier methods. In comparison to the previous study of oncocin^6^, the increased experimental depth has moderately increased assay performance that may allow decreased number of replicate-conditions. The developed parameters (including host transformation efficiency and read depth filter) allow greater precision with fewer replicates and specific (yet fewer) induction levels in identifying active library variants (Supp Figure 4). However, there is very low sensitivity when removing replicates; where again, specific reduced induction levels maintain high sensitivity in identifying most active variants. This suggests that given high experimental depth the assay could be stream-lined, where removing replicate-conditions from the whole may develop stronger stringencies and performance must consistently increase in differences from the wild-type baseline to surpass significance thresholds with these fewer samples. This may allow for expanded sequencing depths of individual experiments and larger libraries able to be screened when searching for the most active antibiotic candidates.

Even with the rich data provided in this study, further inquiries would still be necessary to parse out a more global view of multi-mutational impacts on antibiotic activity. For example, we did not examine alternative locations of PRP-domain inclusion, chimeric exchanges of active segments between peptides, or extensive specificity analysis. Examination and understanding of these phenomena would continue development of these peptides into effective antibiotic candidates.

## Materials and Methods

### Gene construction

Wild-type PrAMP genes (heliocin, pyrrhocoricin, Met, Api, and Tur1a; heliocin with three stop codons after the initial coded residue was used as a negative control) were constructed with overlap assembly of oligonucleotides via PCR. A pET vector was digested using NheI and BamHI restriction enzymes with added calf intestinal phosphatase, and all constructs were gel extracted and purified. The PrAMP gene and vector were ligated using NEBuilder HiFi Assembly and transformed in 10β *E. coli* cells (New England Biolabs). Plasmids were validated by Sanger sequencing.

The *n/n*+1 and *n/n*+2 libraries were constructed with overlapping primers containing degenerate codons NNK at both selected sites for each sub-library. Each sub-library was individually constructed using PCR, gel extracted, and purified. All sub-libraries were combined together as a single mixture for each design. Assembly was done using NEBuilder HiFi Assembly using 0.5 pmol of DNA in a 5:1 ratio of gene:vector. Constructs were purified, concentrated using a MinElute purification column (Qiagen), transformed into electrocompetent 10β *E. coli* cells, and plated on lysogeny broth (LB) plates with 50 µg/mL kanamycin (kan) overnight.

Using a Poisson distribution, it was calculated ∼250x library oversampling achieves at least 50 copies of each library variant present for >99% of variants in each IPTG concentration for each SAMP-Dep replicate (approx. 3 million replicate transformants for Met and 4 million for Api), if equimolar for all variants. To get this level of oversampling for each SAMP-Dep replicate transformation, library DNA was harvested via miniprep (Epoch Life Science) on all transformed and plated cells. All library DNA stocks for SAMP-Dep transformations were above 450 ng/uL for each peptide.

### SAMP-Dep cell growth

Tests for clonal self-depletion were done as in Dejong et al.^6^ Wild-type PrAMP constructs were individually transformed into T7 Express LysY/I^q^ *E. coli* cells. After outgrowth in SOC media, a second outgrowth was done for 1 hr in LB with 50 µg/mL kan. Transformations were then diluted and incubated at 37 ºC with 250 rpm for 12 hours in 0 -0.5 mM IPTG. OD_600_ measurements were taken every 10 minutes with a BioTek Synergy H1 microplate reader.

We used the SAMP-Dep protocol, introduced in Dejong et al.^6^, in a slightly modified form. Previous data demonstrated diminishing benefits to sensitivity and precision beyond three IPTG induction concentrations.^6^ Midpoint concentrations of IPTG were performed for each PrAMP member at 2x serial dilutions from full induction at 0.50 mM (0, 0.016, 0.031, 0.063, 0.13, 0.25, 0.50 mM IPTG) and 0.10 mM steps (0, 0.10, 0.20, 0.30, 0.40, 0.50 mM IPTG) to determine induction levels that offered identification of increased and decreased potency of variants from all wild-type baselines. These concentrations were determined to be 0, 0.15, and 0.50 mM IPTG as uninduced, fully induced, and greatest difference from both at wild-type baseline. Previous data indicated potentially substantial benefits to increasing the number of experimental replicates^6^; thus, the current study utilized quadruplicates. For each replicate, 5 µL of highly concentrated pET-PrAMP DNA was transformed into 50 µL of competent T7 Express LysY/I^q^ *E. coli*. These were recovered in SOC medium for 1 hour. After 1 hour, 10 µL of the 1 mL mixture was serially diluted and plated to determine each library’s transformation efficiency. The remaining 990 µL was pelleted at 6000*g* and suspended in 1 mL LB with 50 µg/mL kan, incubating at 37 ºC shaking for one hour. After antibiotic incubation, each transformation was split into four 250 µL partitions for each IPTG condition and a pre-induction condition. The DNA from this pre-induction condition was immediately harvested via miniprep and stored. 250 µL partitions were added to 9.75 mL of LB with 50 µg/mL kan with each IPTG concentration and incubated at 37 ºC with shaking. After 7.2 hours of incubation, supernatant was decanted, then DNA was harvested and stored. Replicates were performed on four different days.

### Deep sequencing

From each sample (PrAMP/condition/replicate), the same forward and reverse primers were used to amplify the AMP gene variants. Each sample was subsequently amplified with forward and reverse primers with a unique barcode. The resulting purified amplicons were mixed at equimolar amounts (Met and Api pooled together) and sequenced via Illumina NovaSeq 6000 on an SP Flow Cell at the University of Minnesota Genomics Center.

Sequences were filtered via USearch to accept reads with less than .5% expected error rate. Merged sequences were filtered by observations from only uninduced-full incubation sample (0 mM IPTG-7.2 hour incubation), having to pass 20 reads for all replicates to be included in further sequence-function analysis. This read threshold was determined by the calculated number of reads that would have the potential of a maximum depreciation to surpass the wild-type sequence potency (Supp Fig 3).

### Sequence-Function Analysis

For all filtered sequences of each PrAMP, growth rate was calculated as outlined in Dejong *et al*.^6^. *g*_*p*,0mM_ of 12.5 was used as the standard growth rate for an uninduced culture. Negative controls (growth rate:IPTG induction slope of zero) were defined as sequences with a stop codon within the first three codons since they should have no attainable activity. The potency metric was calculated as the growth rate:IPTG induction slope relative to these baselines. The final slope used for analysis was the average of calculated slopes from all four experimental replicates. Significance was calculated through an independent T-test between each variant’s four replicate slopes to the wild-type sequence’s four replicate slopes. All python codes and deep sequencing counts available at: https://github.com/HackelLab-UMN.

### Epistatic Interaction

Epistasis was defined as the change in performance of multimutants beyond the additive properties of each single mutation. This was calculated by the equation:

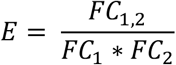

In this equation, FC_1,2_ signifies the fold-change performance of the double mutant in relation to the wild-type. FC_1_ and FC_2_ are defined as the fold-change performance of each single mutant in relation to the wild-type. The log values of the calculated E were plotted against FC_1,2_ for all double mutants. Variants within the bottom left quadrant of the graph and top right were characterized as important since they caused highly deleterious and highly synergistic interactions, respectively. The variants in the top left and bottom right quadrants were not characterized as important since they represent interactions that are not as additive as expected for deleterious and advantageous mutations, respectively.

### Minimal Inhibitory Concentration (MIC) Assay

PrAMP candidate variants were selected for external exposure based on significance of all replicate slopes in relation to all wild-type variant replicate slopes by an independent t-test. For each individual PrAMP tested, three of the top ten performing sequences were selected, with one special case from each PrAMP, the most statistically negative peptide, and the wild-type sequence. These candidates were produced using solid-phase synthesis from ProteoGenix.

Six strains of susceptible species outlined in Lai (2019)^26^ were used as candidates for external exposure of PrAMP variants from SAMP-Dep. The *E. coli* strain T7 LysY was selected as the positive control to directly contrast toward the internal-only exposure within the SAMP-Dep platform. *E. coli* strains JW0013, BW25113, and FVEC 638 were selected based on their properties as a DnaK knockout, widely tested standard, and commensal strain, respectively [Keio Collection, GenoBase, and Minneapolis VA Hospital]. *S. enterica* strains serovar Typhimurium ATCC SL1344 and serovar Enteritidis MH91989 were selected as other common pathogenic species [provided by University of Minnesota]. The MIC assay protocols follow those outlined in DeJong et al. (2021)^6^. Species were diluted and plated on LB-agar. Single colonies were selected for overnight culture in 3 mL of LB media within a dynamic incubator at 37 ºC. On the day of the assay, 10 µL of this overnight outgrowth were inoculated in 3 mL of LB media. These samples grew for 2 hours until they reached exponential phase. This outgrowth was diluted to 5 x 10^5^ CFU/mLand 75 µL was added to 75 µL of 3x serial dilutions of each PrAMP variant in LB based on activity levels previously seen (150, 50, 16.7, and 5.6 µg/mL at final concentration for Met, and 60, 20, 6.7, and 2.2 µg/mL at final concentration for Api) to determine MIC level. OD_600_ measurements were taken every 9 hours with BioTek Synergy H1 microplate reader. Between timepoints, plates were incubated at 37 ºC while shaken at 250 rpm. Samples were conducted in triplicate on two separate days.

To determine MIC, media only growth controls were done for each plate. Minimum inhibitory concentration was defined as the minimum concentration of PrAMP causing a mean OD_600_ measurement below two standard deviations of the mean media only OD_600_ measurements at 9 hours. If differing MIC values were determined on different replicate days, the average was taken. ≥ was used to signify one replicate showed activity at this AMP concentration and the other did not show any measurable inhibition.

## Acknowledgements

This research was supported by the National Institutes of Health (R01 GM121777). We appreciate support from the University of Minnesota Genomics Center.

