## Supplement for "Sequence-function mapping of proline-rich antimicrobial peptides"

Jonathan Collins[1], Adam McConnell[1], Zachary D. Schmitz[2], Benjamin J. Hackel[1,2]
[1]Biomedical Engineering, [2]Chemical Engineering and Materials Science
University of Minnesota, Minneapolis, MN 55455

| AMP | -2 | -1 | 0 | 1 | 2 | 3 | 4 | 5 | 6 | 7 | 8 | 9 | 10 | 11 | 12 | 13 | 14 | 15 | 16 | 17 | 18 | 19 | 20 | 21 | 22 | 23 | 24 | 25 | 26 | 27 | 28 | 29 |
|---|---|---|---|---|---|---|---|---|---|---|---|---|---|---|---|---|---|---|---|---|---|---|---|---|---|---|---|---|---|---|---|---|
| Oncocin | | | | V | D | K | P | P | Y | L | P | R | P | R | P | P | R | R | I | Y | N | R | | | | | | | | | | |
| Apidaecin 1b | G | N | N | R | P | V | Y | I | P | Q | • | • | • | P | H | P | • | L | | | | | | | | | | | | | | |
| Arasin 1 | | | S | R | W | P | S | • | G | R | • | • | • | F | • | G | • | P | K | P | I | F | R | P | R | P | C | | | | | |
| Heliocin | | | R | F | I | H | • | T | • | R | • | P | • | Q | • | R | • | P | V | I | M | • | A | | | | | | | | | |
| Metalnikowin 1 | | | | • | • | • | • | D | • | R | • | • | • | • | • | • | N | M | | | | | | | | | | | | | | |
| Pyrrhocoricin | | | | • | • | • | G | S | • | • | • | • | • | T | • | • | • | P | • | • | • | • | N | | | | | | | | | |
| Tur1a | R | R | I | R | F | R | • | • | • | • | • | • | • | G | R | R | P | • | F | P | P | P | F | P | I | P | R | I | P | R | I | P |

| AMP | Length | % P | % R | Charge | PRPs | PRP Org. |
|---|---|---|---|---|---|---|
| Oncocin | 19 | 32 | 26 | +5 | 1 | overlap |
| Apidaecin 1b | 18 | 33 | 17 | +3 | 1 | non-overlap |
| Arasin 1 | 25 | 36 | 24 | +7 | 2 | non-overlap |
| Heliocin | 21 | 29 | 24 | +5 | 0 | none |
| Metalnikowin 1 | 15 | 33 | 20 | +2 | 1 | overlap |
| Pyrrhocoricin | 20 | 25 | 15 | +3 | 2 | non-overlap |
| Tur1a | 32 | 38 | 31 | +10 | 1 | non-overlap |

**Figure S1** *PrAMPs selected for study*. The first table shows selected PrAMP homology to oncocin, where again residues identical to oncocin are indicated as •. Residues homologous (BLOSUM62 ≥0) to oncocin amino acids are colored gray. Next, chemical and sequence characteristics are displayed. Shading, from green to white to red, displays characteristic similarity to oncocin.

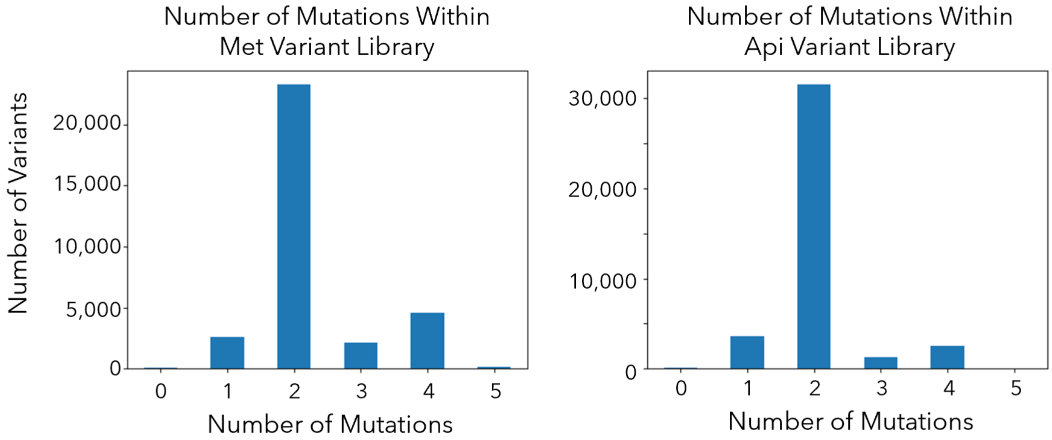


**Figure S2** *PrAMP sequence variants identified within library.* Histograms show the number of sequences screened within the SAMP-Dep assay for each number of mutations present.

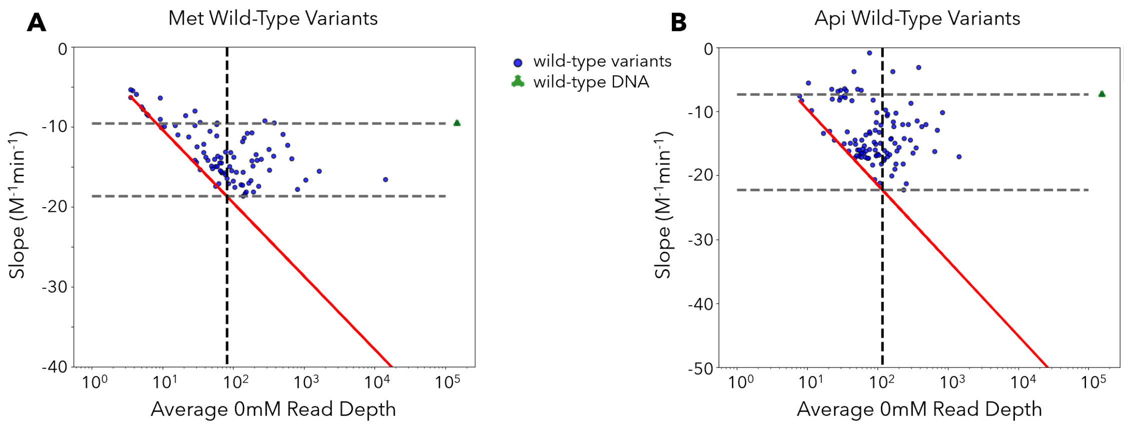


**Figure S3** *Maximum calculable slopes based on average uninduced sample read depth*. Plots show wild-type variants calculated slopes compared to their read depth for A) Met and B) Api. The vertical dashed line indicates the read depth threshold necessary to calculate a slope greater than the maximum wild-type variant slope. The red line indicates the maximum slope able to be calculated based on uninduced read depth. The first horizontal dashed gray line shows the wild-type DNA slope and the second horizontal dashed gray line shows the maximum wild-type variant slope.

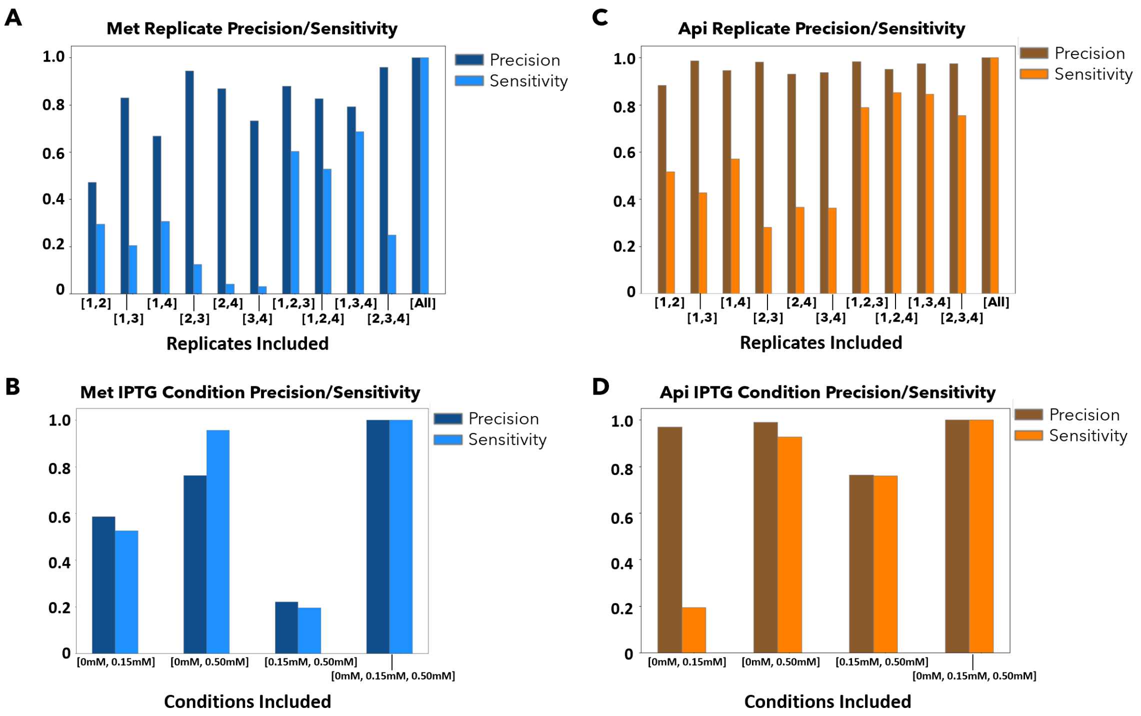


**Figure S4** *Sensitivity and Precision of SAMP-Dep.* Bar plots show the individual precision and sensitivity as calculated in Dejong et al. (2021)[41] based on A) Met replicates, B) Api replicates, C) Met induction concentrations, and D) Api induction concentrations.
